# An integrated synthetic biology and robotics approach for neutralising landmines in post-war communities

**DOI:** 10.64898/2026.01.20.700574

**Authors:** Y. Basti, S. Williams, E. Aellen, F. Muci, I. Amri, A. Davila, A. Schlüter, A. Dao, P. Meyer, J. Dembska, R. C. Smith, B. D. McCabe

**Affiliations:** EPFL, Lausanne, 1015, Switzerland; 1nstitute of Materials, EPFL, Lausanne, 1015; 1nstitute of Bioengineering, EPFL, Lausanne, 1015; Brain Mind Institute, EPFL, Lausanne, 1015

**Keywords:** TNT biodegradation, bioremediation, unexploded ordnance (UXO), drone, synthetic biology

## Abstract

Unexploded ordnances (UX.Os) and landmines endanger lives and hinder the economic progress of communities living in post-conflict zones. Currently, the primary method for clearing UX.Os relies on metal detection and manual removal of UX.Os - an expensive, time-consuming, and hazardous process. This study, derived from the 2024 EPFL iGEM project SYNPLODE, presents a new approach that integrates synthetic biology and aerial drone robotics, proposing a novel, end-to-end, safe, and efficient solution to address UX.Os. Starting from bacteria engineered to detect and degrade 2,4,6-trinitrotoluene (TNT), a common explosive in landmines, our solution is designed for three main tasks: detecting **TNT** and RDX, breaking these compounds down into non-explosive byproducts, and confirming explosive neutralisation. To deploy this solution safely in UXO-contaminated areas, we designed, built, and tested an aerial drone capable of spraying explosive-degrading bacteria. Combining synthetic biology, robotics, mathematical modelling, and affected community engagement, our solution aims to improve UXO and landmine clearance by offering a scalable and cost-effective approach for deactivating UX.Os without risking human lives.

## Introduction

Unexploded ordnances (UX.Os) and landmines are persistent remnants of war that pose significant threats to human life, impede economic growth, and restrict land use long after hostilities have ceased. Many communities live exposed to these risks, but traditional demining methods are hazardous, time-consuming, and costly [1].

Bioremediation offers a promising alternative for addressing explosive contamination. Previous studies have successfully demonstrated the use of engineered biological com ponents as biomarkers for detecting explosive compounds[2, 3]. Moreover, genetically modified bacteria have been shown to accelerate the degradation of certain explosive molecules beyond their natural decay rate [4, 5]. Building on these findings, our 2024 iGEM project, SYNPLODE1^1^ aimed to develop a comprehensive solution for landmine clearance and enhance the overall efficiency of demining operations. By integrating synthetic biology, community engagement, and robotics, we designed bacterial vectors capable of degrading explosives within landmines, supported by a mathematical model to predict their performance. To deploy these bacteria effectively, we developed an aerial drone system tailored for minefield environments. Throughout the project, we engaged with a range of stakeholders-including deminers, non-governmental organizations (NGOs), and academic researchers-to better understand real-world challenges, identify potential limitations, and ensure the practicality and safety of our approach.

Our objective was to target two of the most prevalent explosive compounds found in landmines and unexploded ordnance (UX.Os): 2,4,6-trinitrotoluene (TNT) and 1,3,5-trinitro-1,3,5-triazinane (RDX) [3]. We engineered bacterial systems capable of detecting these compounds, degrading them into non-explosive derivatives, and signaling the successful completion of this process.

To achieve TNT biodegradation, we designed a system to overexpress the enzymes NemA, NfsA, and NfsB from *E. coli* strain K12. These enzymes reduce TNT’s nitro groups to amino groups, producing non-explosive products such as 2,4-diamino-6-nitrotoluene [6–8], 2-amino-4,6-dinitrotoluene and 4-amino-2,6-dinitrotoluene (ADNT) [9], and 2,4,6-triaminotoluene (TAT) [6]. Notably, NemA alone has been shown to triple TNT degradation rates [4] and can also degrade other explosives such as glycerol trinitrate (GTN) and pentaerythritol tetranitrate (PETN) [10]. NfsA and NfsB, both nitroreductases, also play key roles in this process [10]. Their combined overexpression is thus expected to further enhance TNT breakdown. Importantly, the degradation process releases ammonium, providing a usable nitrogen source that supports bacterial growth in TNT-contaminated environments [4].

For RDX degradation, we transferred the *Rhodococcus rhodochrous* enzymes XplA and XplB into *E. coli* for heterologous expression [5]. These enzymes degrade RDX through denitration and ring cleavage, producing byproducts such as nitrite, formaldehyde, nitrous oxide, carbon dioxide, and formate [5, 11]. Like TNT, RDX also serves as a nitrogen source for the degrading organism [5].

To verify complete degradation of both TNT and RDX, we designed a visible output system using a 3-input genetic AND gate. This circuit produces a detectable signal only when three specific conditions are simultaneously met: the absence of both TNT and RDX, and the presence of degradation byproducts [12]. This ensures the signal only appears when clearance is achieved.

To address biosafety, we developed a kill-switch mechanism to ensure bacterial death in the absence of target compounds. Based on the CcdB/CcdA toxin-antitoxin system [13, 14], our design permits bacterial survival only in the presence of TNT or RDX. Once these compounds are degraded, CcdB is expressed, triggering cell death.

A key challenge in applying this system is delivering bacteria to the explosive materials, which are typically encased in metal, plastic, rubber, or wood shells (Fig. 1) that can block bacterial entry [15].

**Figure 1.**
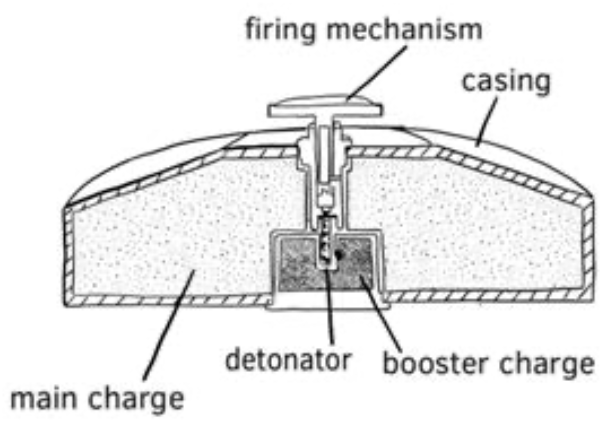
Schematic of a landmine. The explosive load is encapsulated in a casing. Fig. adapted from [16].

To investigate practical delivery methods, we consulted with demining experts from DanChurchAid (DCA) and the Health and Social Care Organization in Iraq (IHSCO). They confirmed that, over time, metal landmine casings buried underground can develop cracks due to environmental exposure. These cracks allow water infiltration and leaching of explosive chemicals into the surrounding soil. The degradation rate of the casing depends on factors such as humidity, temperature, and sun exposure. Many UXOs remain buried long enough for such degradation to occur, creating entry points for a bacterial solution.

Based on this insight, we developed a drone capable of flying over minefields and spraying the degrading solution directly onto landmines. This approach enables remote deployment, enhancing safety for deminers. The method allows for targeted delivery without the need to saturate the entire area.

To further automate the process, the drone can be paired with detection systems. One common method currently in use involves drones equipped with infrared cameras to identify landmines based on thermal signatures that differ from the surrounding soil [17]. However, this technique is limited by the need for a sufficiently large temperature contrast. As an alternative, we implemented biosensors that emit visible light in response to TNT and DNT vapors [2, 3], which can be detected with standard cameras. By spraying the detection solution across the minefield, we can identify both the locations of buried landmines and areas where explosive chemicals have leaked into the soil. This dual detection capability supports both demining and remediation of explosive-contaminated environments. Accordingly, we equipped our drone with a standard camera to test the feasibility of using biosensors for visible signal detection.

## Methods

### Stakeholders interviews

To understand the landscape of current demining efforts and evaluate the potential of integrating a novel synthetic biology approach into existing workflows, we interviewed a diverse set of stakeholders involved in demining operations. We received advice and information from six NGOs, two defence specialists, two academic researchers, and two public organisations representatives, all from different parts of the world with various roles or expertise in the field of demining (exhaustive list in the Supplementary Materials). Each interview lasted between 45 and 60 minutes and was conducted virtually or in person. The questions were tailored to each participant and focused on the current state and limitations of demining methods, regulatory challenges, and the potential strengths and weaknesses of our solution. These discussions helped us gain insight into current demining practices, operational challenges, and the perceived risks and benefits of our proposed synthetic biology-based intervention in the field.

### Modelling of detection process

In our demining strategy, we propose a detection system based on bacterial luminescence in response to TNT vapours escaping from buried landmines. To assess the feasibility of such a method, we developed a simulation model to predict the diffusion of explosive compounds–particularly TNT–through soil. As the model is geared towards eventual field implementation, it is designed with a minimal number of parameters to allow for fast calibration in real-world conditions. Therefore, we adopt a simplified approach by modelling gas transport in soil using Fick’s Law [18] :

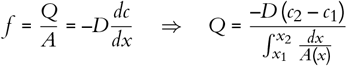

Where *f* is the flux density, *Q* is the flux, *A* is the area, *D* is the diffusion coefficient, *c* is the concentration and *c*_1_ and *c*_2_ are the concentrations at positions *x*_1_ and *x*_2_. We define the conductance *K* as:

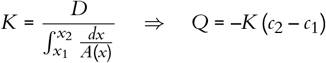

In view of a numerical implementation, we discretise the system using a simple lattice grid, which reduces the model to three parameters: the average soil conductance *K*µ, the variance in conductance 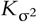, to account for soil heterogeneity, and the vapour leakage rate *u* from a mine.

### Synthetic biology

To demonstrate the potential of synthetic biology as a viable solution for landmine clearance, we needed to engineer bacteria with improved ability to degrade explosives while integrating essential safety measures for environmental release. Furthermore, techniques were required to measure the bacteria’s efficiency in degrading TNT and RDX.

### Plasmid design ^2^

#### TNT degradation and detection

To engineer *E. coli* for enhanced TNT degradation, we designed a plasmid to overexpress the TNT-degrading enzymes NemA, NfsA, and NfsB (Fig. 2a). These enzymes were cloned downstream of the yqjF3rd promoter, an improved version of the yqjF TNT-sensitive promoter, developed and characterised by the 2020 iGEM NEFU team [2]. The wildtype yqjF promoter activates gene expression in the presence of TNT or its breakdown products such as DNT, and/or THT [3, 19–21]. To enhance promoter sensitivity, a second plasmid included the yhaJ1st variant of the yhaJ transcriptional activator [2] under the constitutive J23100 promoter.

**Figure 2.**
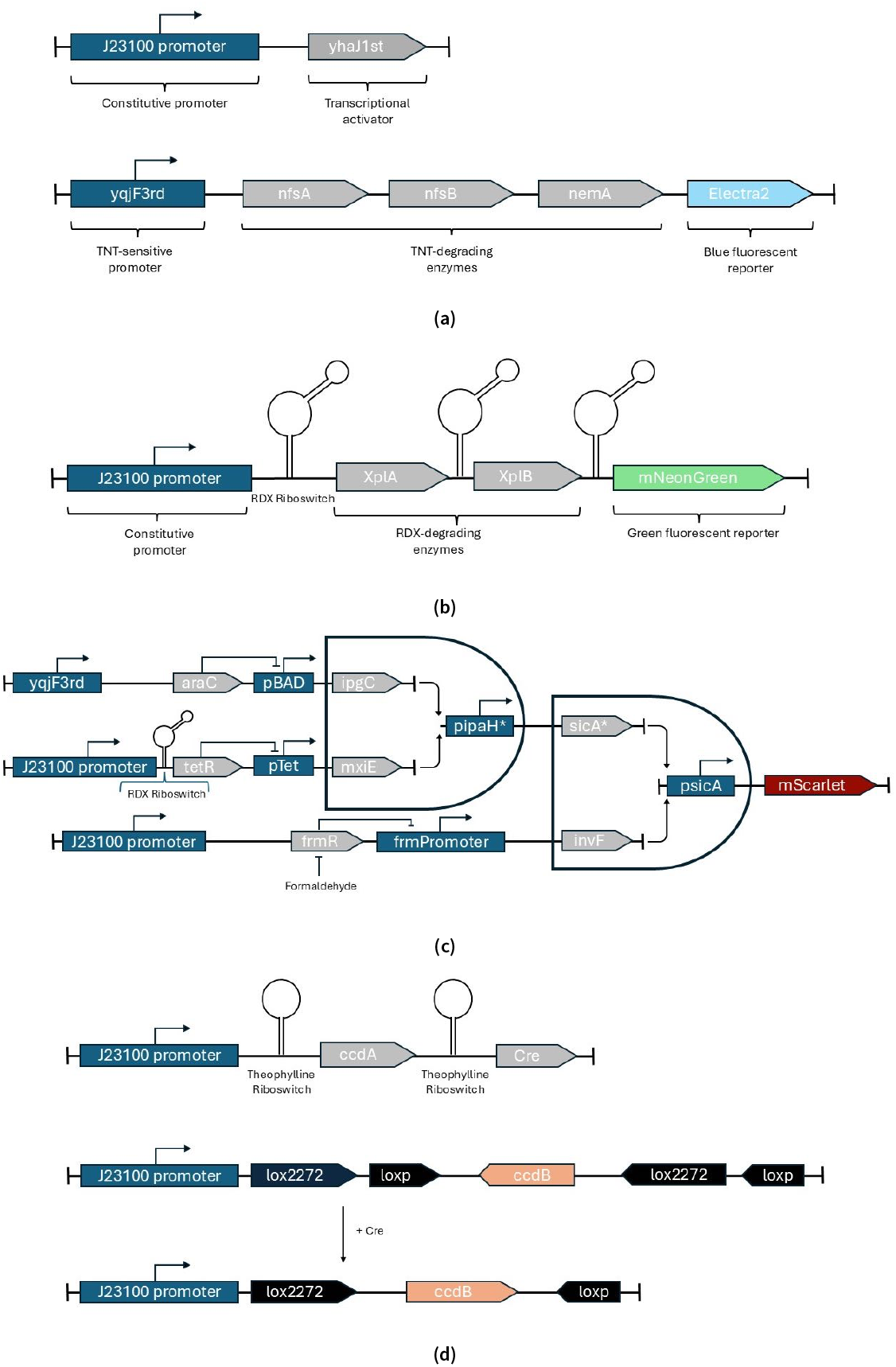
Bacterial engineering. **(a)** Genetic constructs for TNT detection and degradation. The engineered yhaJ1st transcriptional activator of the yqjF3rd promoter is constitutively expressed under the control of the J23100 promoter, while the TNT-inducible promoter yqjF3rd triggers the expression of TNT-degrading enzymes nfsA, nfsB, and NemA, along with the blue reporter fluorescent protein Electra2. **(b)** Genetic construct for RDX detection and degradation. An RDX-sensitive riboswitch induces the expression of RDX-degrading enzymes XplA and XplB, along with the green fluorescent protein mNeonGreen. **(c)** Implementation of the genetic logic gate. In the presence of TNT, yqjF3rd is active, triggering the expression of araC, which represses the pBad promoter. Similarly, RDX presence leads to the expression of tetR, which represses the pTet promoter. FrmR is produced constitutively to repress the frm promoter, but is inhibited by formaldehyde. **(d)** Theophylline-dependent activation of the kill-switch using a FLEx switch. The CcdB gene is cloned in antisense orientation relative to the J23100 promoter and flanked by loxP and lox2272 recognition sites. Theophylline induces Cre recombinase expression, which excises and inverts the CcdB sequence, leading to constitutive toxin production.

**Figure 3.**
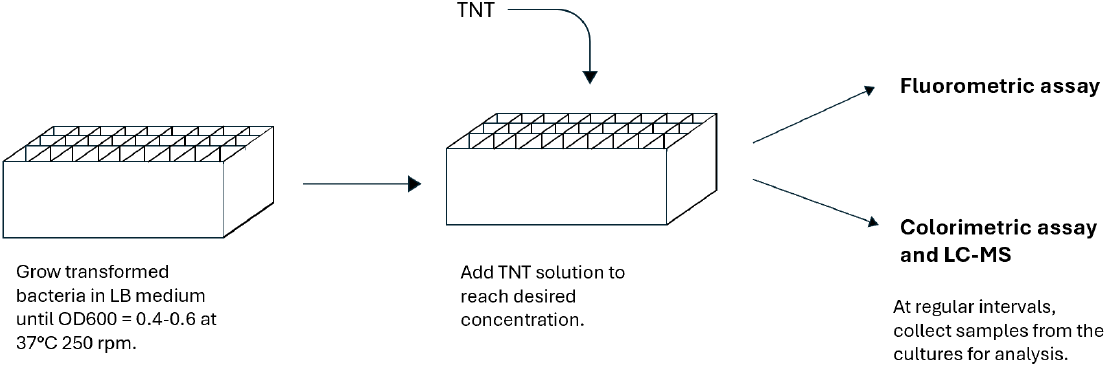
TNT degradation experiments : Deep-well plate containing the transformed bacteria incubated in LB medium to reach the optimal OD. Various amounts of TNT are added to the bacterial solution. The mixture is then analyzed using a FLuorometric assay, a Colorimetric assay and LC-MS at regular time intervals.

**Figure 4.**
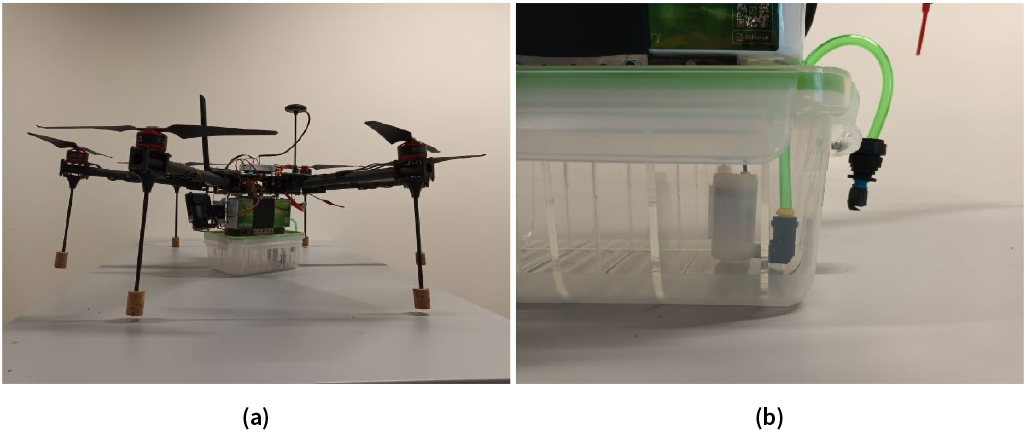
Our hexacopter drone (a), with the tank, pump, pipe and nozzle of the spraying system (b).

The first construct included the Electra2 blue fluorescent protein [22] to visually indicate the presence of TNT. Sequences were then incorporated into pGEX-4T-1-H vectors from GenScript with kanamycin and ampicillin resistance markers.

RDX degradation and detection

RDX-degrading enzymes XplA and XplB were cloned downstream of a RDX-sensitive riboswitch, together with the mNeonGreen fluorescent protein, ensuring translation in response to RDX binding [23, 24] (Fig. 2b). The polycistronic construct was constitutively transcribed under the control of the J23100 promoter. The constructs were incorporated into the pCDFDuet-1 backbone from GenScript.

#### AND gate design

We designed a 3-input AND gate (adapted from [12]) using the TNT-sensitive yqjF3rd promoter and the RDX-activated riboswitch to regulate transcriptional repressors (Fig. 2c). In our design, these repressors inhibit the gate until TNT and RDX are fully degraded. As third input, we used the formaldehyde-inducible frm promoter, activated by formaldehyde [25], a by-product of RDX degradation. FrmR represses the frm promoter unless formaldehyde is present. All three inputs are required for activation, signalling the absence of TNT and RDX and the presence of degradation by-products. For initial tests, we planned to use mScarlet fluorescent protein [26] as a marker.

#### Kill-Switch design

The kill-switch mechanism was conceptualized to express the CcdB toxin under the control of the constitutive J23100 promoter, while coupling the CcdA antitoxin production to the TNT-activated yqjF3rd promoter and the RDX-sensitive riboswitch. This design allows cell survival only in the presence of TNT or RDX, triggering cell death once the target compounds are depleted. To provide conditional kill-switch activation upon deployment, we planned to use a Lox FLEx (flip-excision) switch [27] to activate CcdB expression before deployment (Fig. 2d). We cloned the CcdB gene in antisense orientation relative to its promoter and flanked it by loxP and lox2272 recognition sites. Cre recombinase, expressed under the control of a Theophylline riboswitch [28], enables conditional CcdB expression through sequence inversion. This switch ensures that TNT and RDX dependency for cell survival would only activate once the culture is ready for deployment on a minefield or other contaminated area. In this solution, Theophylline application would be required immediately prior to the deployment of the bacteria. To account for the time between the bacteria reaching the landmine and penetrating its casing, we incorporated a CcdA gene to the Theophylline-controlled system, which drives Cre expression. This allows the bacteria to survive for a few hours before reaching the target and relying on TNT and RDX for survival.

#### Bacterial Culture and TNT Fluorometric Assay

*E. coli* BL21 cells, transformed with TNT-degradation and detection plasmids, were cultured in Lysogeny Broth (LB) medium supplemented with kanamycin and ampicillin(Sigma Aldrich and Adipogen AG) and incubated at 37°C with shaking at 250 rpm. Bacterial growth was monitored using optical density at 600 nm (OD600), and cells in the logarithmic growth phase (OD600 = 0.4–0.8) were used. For experiments, bacterial cultures were diluted 100x into fresh LB medium with varying TNT concentrations. Negative controls included LB without TNT, TNT in LB without bacteria, and wild-type BL21 with TNT. We conducted experiments over 50 hours. Aliquots (100 µL) were transferred to flat-bottom 96-well plates (ThermoFisher) for measuring relative fluorescence units (RFU) and OD600 (with Cytation 5,BioTeK). Cultures were incubated at 37°C between measurements with consistent shaking (250 rpm). For the calibration 100 µL LB medium were used and the fluorescence was measured with excitation at 403 nm and 454 nm emission wavelength.

#### Dicyclohexylamine TNT Colorimetric Assay

This assay quantified TNT concentrations and evaluated bacterial degradation efficiency using the interaction of TNT with dicyclohexylamine, which forms a red-violet charge-transfer complex [29]. The intensity of the complex correlates with TNT concentration. A TNT stock solution of 1000 µg/mL (Armasuisse and Agilent) was prepared with acetone and 0.05% (w/v) methyl paraben. A 10:1 (v/v) mixture of dicyclohexylamine (ThermoFisher) and isobutyl methyl ketone (from Chemie Brunschwig AG) (DCHA-IBMK) was used for extractions. This method was adapted for small volumes from Üzer et al. 2005 [29]. 10 µL of 0.05% methyl paraben solution were added to 50 µL of a TNT solution (1:1 acetone and LB medium). After adding 60 µL of the DCHA-IBMK mixture and vortexing, the tubes were centrifuged to separate liquid phases. The red-violet complex formed in the organic top layer was measured for semi-quantitative analysis.

#### Liquid Chromatography–Mass Spectrometry (LC-MS) for TNT Detection

After performing the colorimetric assay, we conducted LC-MS to obtain a precise quantification of the TNT present. Samples from the colorimetric assay were used, with bacteria removed by centrifugation to collect the supernatant. Analysis, conducted at EPFL’s Central Environmental Laboratory (CEL), measured the TNT concentrations over 0 to 12 hours. The Acquity Binary Solvent Manager pump used a mobile phase of MQ water with 2 mM ammonium acetate (Phase A) and methanol with 2 mM ammonium acetate (Phase B). The gradient program transitioned from 90% Phase A (0.300 mL/min) to 90% Phase B over 10 minutes, reverting to initial conditions by 15 minutes. Injection was performed with a 2777 Sample Manager using a 100 µL syringe (10 µL injection volume). Washing protocols included a strong wash (MQ:ACN:MeOH:IPROH, 1:1:1:1) and a weak wash (MQ:ACN, 1:1). Separation employed an Acquity UPLC BEH Phenyl column (1.7 µm, 2.1 x 50 mm), and detection was via the XEVO TQ MS system, which monitored specific retention times and ion transitions. This ensured accurate and reproducible TNT quantification.

### Delivery method

Spraying drones are currently used in agriculture and many efficient models that could deliver a degrading solution on landmines exist. Nevertheless, they are very expensive (DJI Agras T40 - $20’000), and communities impacted by UXOs may not have the resources for such expensive tools. Therefore, we designed our own prototype to both prove the feasibility of this delivery method and test a low-cost alternative to existing agricultural drones, which any user can independently build, repair, and adapt to their specific needs.

We designed a 1 metre diameter hexacopter capable of carrying a payload of 2 litres. Compared to quadcopters or octocopters, hexacopters (6 motors) offer a compromise between minimal thrust required to lift the payload and the cost of having more motors. We chose low KV BLDC motors (Racestar 41x14 mm 400 KV), which spin slowly but generate a high torque, with [15x5.5 inches EOLO carbon fiber propellers](https://sunnyskyusa.com/products/eolo-cn15-5-5-prop), to favor flight stability and endurance over agility. The motors are controlled [40A BLHeli ESCs](https://emaxmodel.com/products/blheli-series-40a-ubec) by and powered with a Swaytronic LiPo battery 6S 22.2V 16000mAh 35C (1.9 kg). We started the assembly with a ZD850 carbon fiber frame, but we had to modify the landing gear – increasing the number of feet from 4 to 6 and lowering the center of gravity – to enable take-offs and landings in versatile conditions such as on rough or inclined surfaces. The tank of the spraying system and its liquid content represent a significant part of the drone mass and have a huge impact on its balance. To prevent disturbances from the liquid flowing back and forth, we laser cut a grid out of PMMA plaques and inserted it to partition our polypropylene tank. The spraying system consists of a small 5V submersible pump, which expels liquid through a 6 mm polypropylene pipe terminated with a nozzle. We equipped the drone with a [Pixhawk flight controller](https://docs.px4.io/main/en/flight_controller/pixhawk.html), a GPS, a telemetry module, an RC receiver, and a GoPro Hero 5 camera, articulated on a two axis gimbal, for first person view (FPV) flights. For testing, we used the PX4 autopilot in combination with the ground station [QGroundcontrol](https://qgroundcontrol.com/) or a 10 channels RC controller. More details can be found on the website of our project [SYNPLODE](https://2024.igem.wiki/epfl/hardware).

### Bioluminescence detection

Finally, to investigate the detection of bioluminescence using a drone camera, we conducted preliminary tests in laboratory conditions using recombinant luciferase proteins (Promega) spread on soil samples. The emitted light was captured in complete darkness with a standard webcam at a distance of 30 cm.

### Data analysis

The data of the TNT concentration over time obtained from the LC-MS measurements and the expression of fluorescent markers under the control of the TNT-responsive yqjF3rd promoter were analysed using R version 4.3.2 in RStudio version 2023.12.0+369. To obtain an OD-normalized version of the LC-MS data to control for variation due to varying abundance of bacteria in the different conditions, we divided the concentration of TNT of each condition at a each time-point by the OD600 values of the respective condition at each time point. For representative purpose we show the un-normalized concentration next to the LB+TNT control condition, which cannot be OD-normalized due to the absence of bacteria, in the main text.

The fluorescence data was OD-normalized by dividing the RFI values of each condition at each time point by the OD600 values of the respective condition at that given time point. Subsequently, the background fluorescence was removed by dividing the OD-normalized RFI values of each condition at each time point by the mean RFI value of the LB+TNT control at that given time point.

## Results

### Modelling

To validate our model, we compared the simulation results to experimental data from [30]. In their study, TNT concentration was measured as a function of depth in a tank setup with a 14 cm internal diameter and a height of 55 cm, filled with 7.5 kg of sand. The TNT source was a 0.5 g crystalline sample placed 5 cm below the surface of the sand.

Currently, engineered bacterial strains designed to detect TNT have a detection limit of approximately 0.04 mg/cm^3^ [31]. To assess their potential for field deployment, we simulated the TNT concentration profile above a landmine buried 5 cm beneath the ground. Figure 5b shows the TNT concentration in an 8 x 8 cm patch of land directly above the mine. Our results indicate that these bacteria can signal the presence of TNT within a 4 cm radius of the mine.

**Figure 5.**
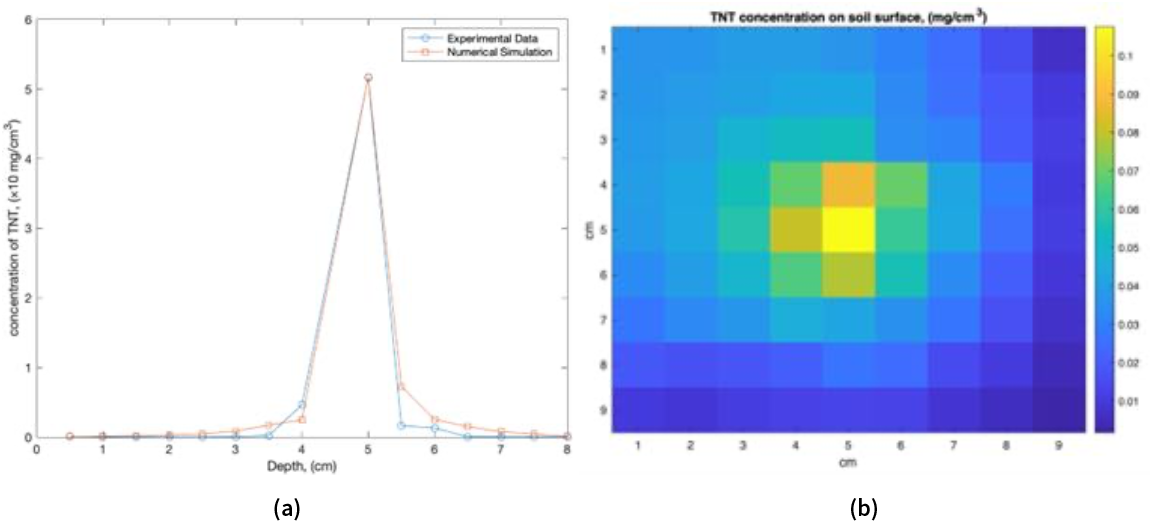
(a) TNT concentration in function of depth. Comparison of simulation results (fitted parameters: 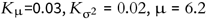) with experimental data [30] (b) Predicted TNT concentration on the soil surface above a landmine buried 5 cm deep.

### TNT colorimetric assay and calibration curve to estimate TNT concentration

To validate the TNT colorimetric assay, we tested a range of TNT concentrations: 70, 50, 40, 5, 0.25, and 0 ng/µL (blank). A decrease in TNT concentration resulted in reduced intensity of the violet dye in the organic phase (Fig. 6a). To assess TNT degradation, bacteria containing the degradation construct were exposed to 30 ng/µL TNT and incubated for 12 hours. In the control sample (sample 1), in the absence of the TNT-degrading bacteria, the violet dye persisted (Fig. 6b). In contrast, no dye was visible in the sample with modified bacteria (sample 2) (Fig. 6b). Native *E. coli* exposed to 30 ng/µL of TNT also show absence of the violet dye (Fig. 6c). To analyse the colorimetric assay results, we used a calibration curve (Fig. 6d) to estimate the unknown TNT concentrations in our samples. The resulting calibration curve is described by the equation: *y* = 6 × 10^−7^*x* +6 × 10 ^−5^, with an *R*^2^ value of 0.8735.

**Figure 6.**
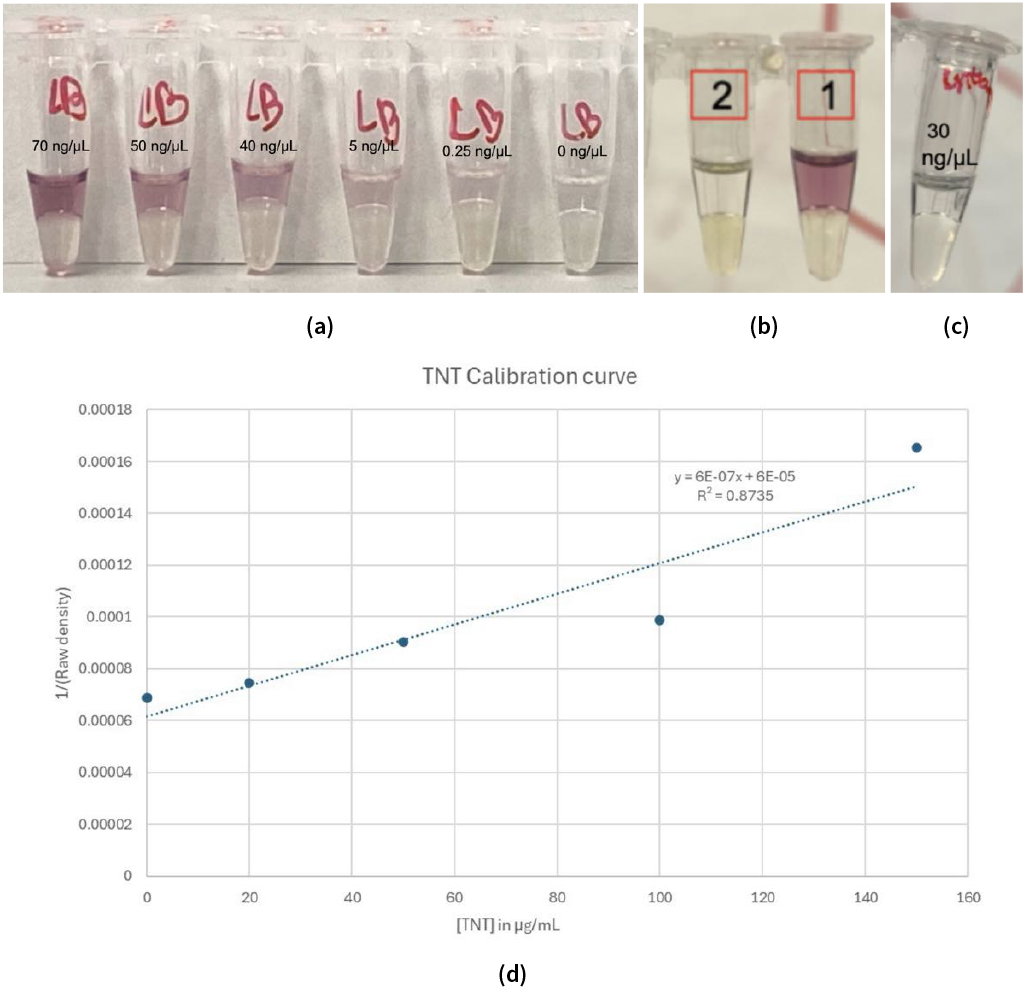
**(a)** TNT Colorimetric Assay: Relationship between TNT concentration (ranging from 70 to 0 ng/µL) and color intensity in the organic phase as observed in the colorimetric assay. **(b)** Comparison of TNT-degrading bacteria with TNT (Sample 2) vs. TNT only (Sample 1): Both samples contain 30 µg/mL of TNT. Sample 2 includes TNT-degrading bacteria, where no purple dye is observed, indicating TNT degradation, while Sample 1 (without bacteria) retains a purple dye, signifying the presence of TNT. **(c)** Wild-Type Bacteria with TNT: Results for Wild-Type *E. coli* BL21 exposed to 30 µg/mL of TNT, showing no purple dye, indicating TNT degradation. **(d)** TNT Calibration Curve: Calibration curve showing quantified TNT degradation using the colorimetric assay.

### Formation of the TNT Meisenheimer complex showing potential presence of a TNT degradation product

When TNT-degrading and wildtype bacteria were exposed to higher TNT concentrations (70 µg / ml), a dark orange product was observed in the different samples (Fig. 7).

**Figure 7.**
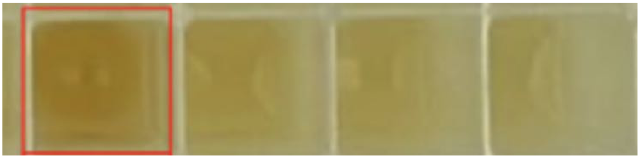
Deep well plate showing a range of decreasing TNT concentrations (from left to right: 70 µg/mL, 30 µg/mL, 0.5 µg/mL, and 0 µg/mL), all incubated with TNT-degrading bacteria. The well highlighted in red represents TNT at a concentration of 70 µg/mL in the presence of TNT-degrading bacteria. A dark orange product is prominently visible in this well.

### LC-MS to quantify TNT concentration

LC-MS results showed significant decrease in the concentration of TNT for both samples containing 70 µg/mL TNT and the engineered bacteria but also for the native *E. coli* in presence of 70 µg/mL. No TNT was observed after approximately 12 hours of incubation (Fig. 8). In contrast, control samples containing only TNT in LB medium without bacteria retained consistent concentrations. However, both bacterial strains exhibited rapid TNT uptake, with initial measurements showing a reduction from 70 to 30 µg/mL within 5 minutes. To enhance degradation rates, we switched from the TNT-inducible yqjF3rd promoter to the constitutive J23100 promoter. Constitutively expressed nfsA showed the fastest decrease in the concentration of TNT during the early stages (0–2 hours).

**Figure 8.**
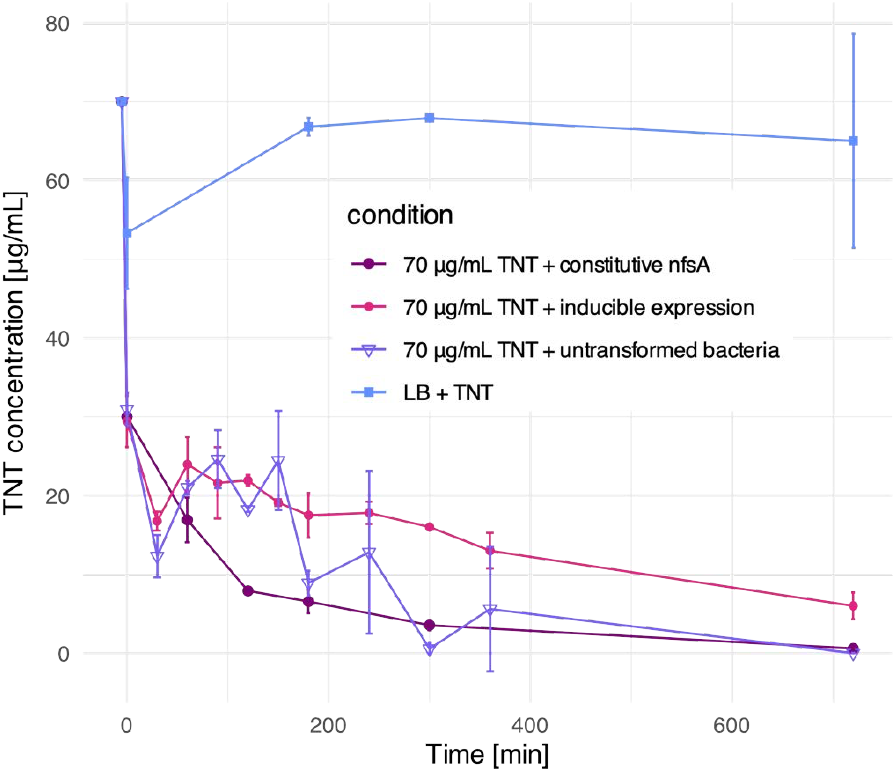
TNT concentration over time with 70 µg/mL input concentration. The purple curve represents the trend line for TNT degradation by bacteria with the constitutive expression of nfsA, showing the steepest slope. The blue-purple and pink curves represent TNT degradation by the bacteria expressing nfsA, nfsB, and nemA under the control of the yqjF3rd promoter and by untransformed E. coli BL21 bacteria, respectively. The blue curve represents the control containing only 70 µg/mL of TNT and LB, showing no decrease of the TNT concentration over time.

**Figure 9.**
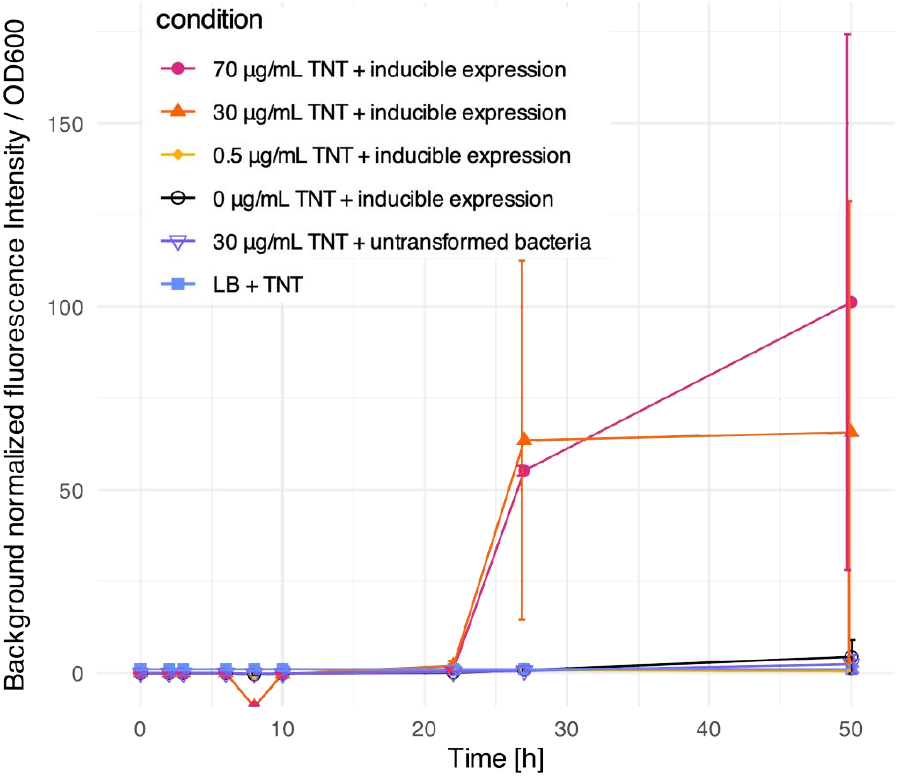
Measurements of Electra2 fluorescence over time for di”erent TNT concentrations. The fluorescence intensity of the pink curve (70 µg/mL TNT) and the orange curve (30 µg/mL TNT), both containing the engineered bacteria, increases affer approximately 20 hours. The blue curve represents the control (LB+ 70 µg/mL TNT) and stays constant over time.

### TNT fluorometric assay to assess the activity of the yqjF3rd TNT sensitive promoter

To investigate the activity of the yqjF3rd promoter in response of TNT, we conducted a fluorometric assay. Activity of the Electra2 fluorescent protein downstream of the TNT degradation constructs was monitored. No fluorescence was observed during the first 12 hours. However, fluorescence was detected after 22 hours in samples exposed to 70 and 30 µg/mL of TNT.

### Delivery method

The drone we built can fly autonomously for 20 minutes carrying a payload of 2 litres. The total cost of the drone is less than 1700 USD. Although we did not test the recognition of contaminated areas using activated TNT biosensors, the FPV system enables a pilot to localise a target like an UXO. The machine is stable enough to spray the solution with an accuracy of 10 x 10 centimetres. Arguably, the payload capacity of our low-cost device may seem small compared to the number of landmines to spray, but its ease of fabrication and set-up may help to scale up the process of demining. Finally, the spraying accuracy demonstrates the feasibility of using a drone to deliver the degrading solution on UXOs or contaminated areas.

### Bioluminescence detection

Having constructed a functional drone, we then sought to investigate the potential of bioluminescent reporters to detect landmines. The luminescent signal on soil samples was of low intensity, but detectable (Fig. 10). With a standard webcam, we were able to extract the localisation of luminescent areas through successive normalisation, thresholding, and morphological operations. Further experimentation is required with the final strain of bacteria to determine the duration of luminescence, its strength, and its sensitivity to explosive chemicals. Such results will decide if bioluminescence can be detected from a drone, for how long and at which distance, and finally if it can be successfully implemented for landmine detection. While the initial constructs utilise fluorescent proteins, it is important to note that they could be readily adapted for field detection by substituting them with luciferases to generate visible luminescence. For example, Electra2 can be replaced with Cypridina luciferase, mNeonGreen with Green Renilla luciferase, and mScarlet with LuciolaRed luciferase [32].

**Figure 10.**
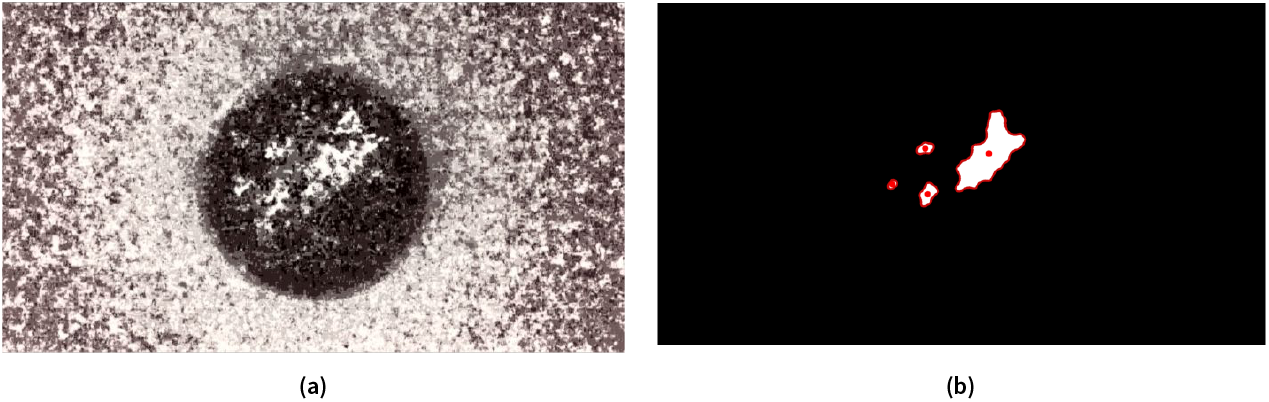
Detection and processing of the signal emitted by luciferase spread on earth in a Petri dish: (a) normalized signal, (b) segmentation of the glowing areas by thresholding and morphological processing.

**Figure 11.**
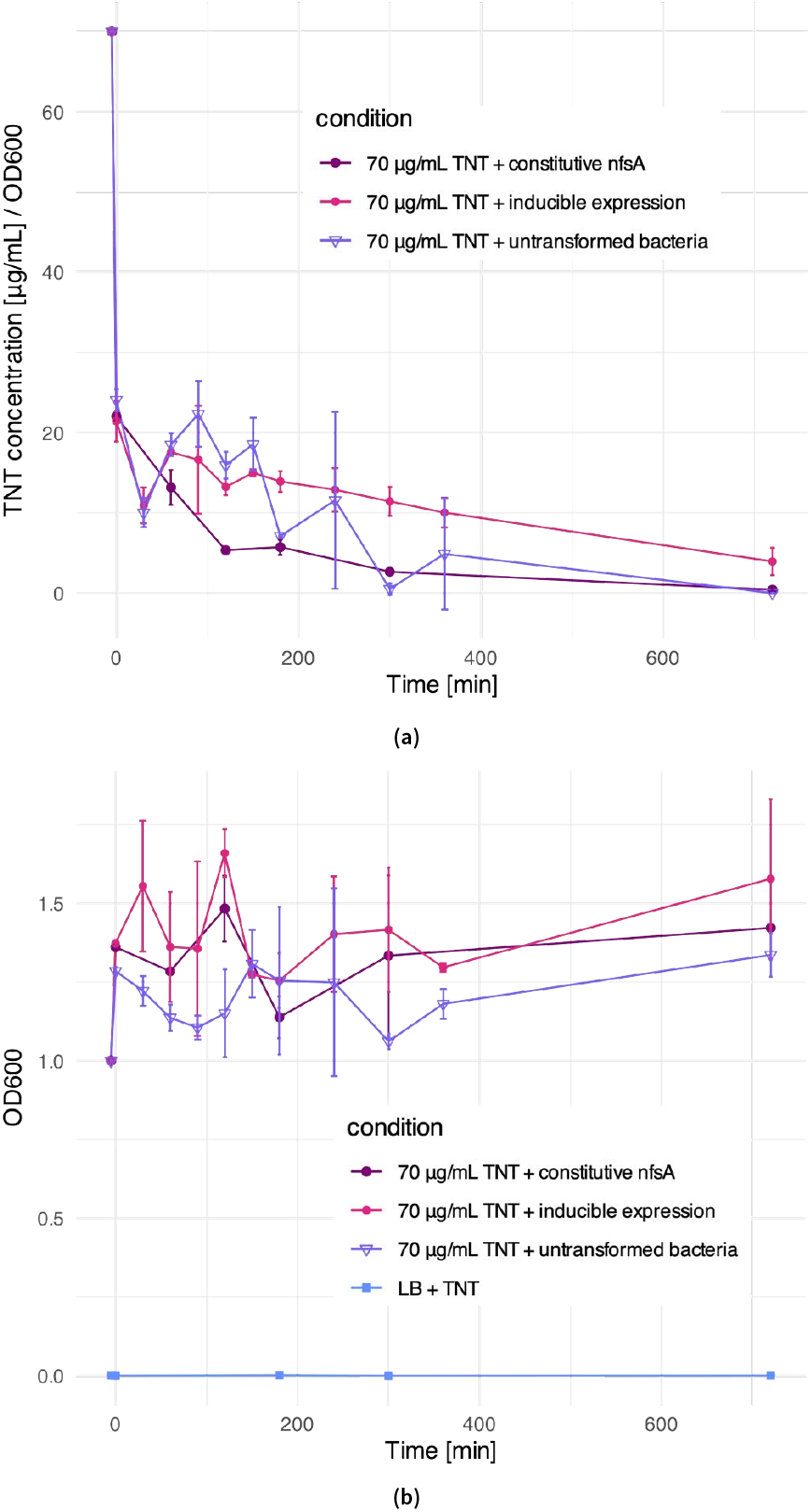
(a) OD-normalized TNT concentration over time with µ0 µg/mL input concentration. The purple curve represents the trend line for

**Figure 12.**
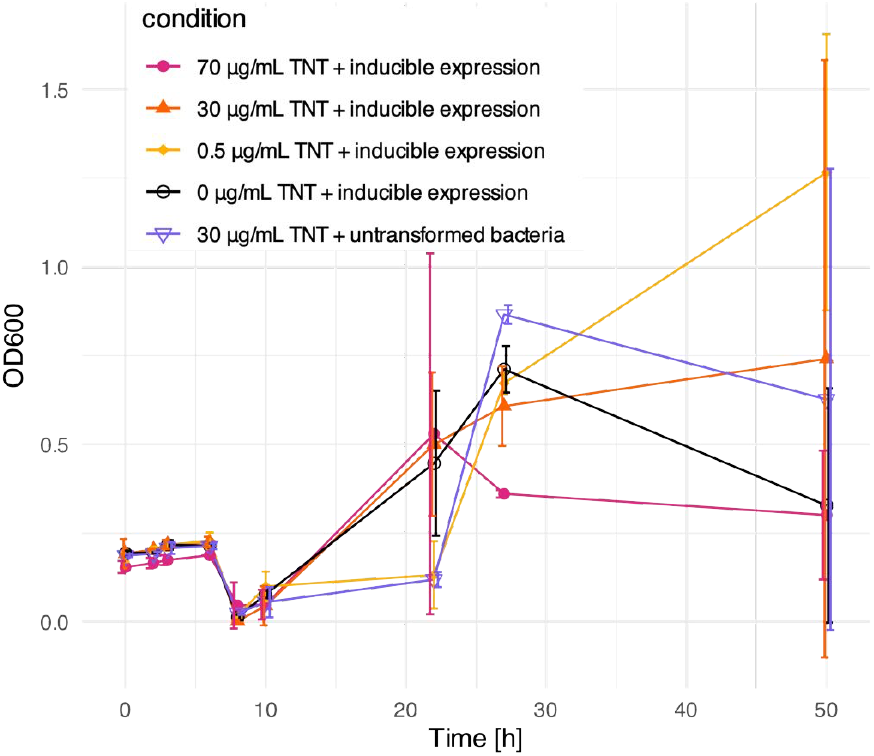
Measurements of OD600 over time for di”erent TNT concentrations in the fluorescence induction experiment.

### Stakeholder Interviews

Engaging with stakeholders provided critical insights into current demining practices and revealed how our synthetic biology-based solution could address key limitations in existing approaches.

Despite technological advances and the use of trained animals such as dogs and rats, stakeholders consistently noted that the core method of landmine clearance remains largely unchanged: “essentially the same technique used over 100 years ago,” according to Bill Morse, chairman of Cambodian Self Help Demining (CSHD). All NGOs we interviewed confirmed that manual clearance remains the dominant method, with modern technologies often being prohibitively expensive and difficult to implement in the field. As a result, demining remains a slow, costly, and dangerous process. Clearing a single minefield can take up to 12 months, with an estimated cost of up to $1,000 per landmine.

Environmental contamination emerged as another pressing concern. Unstable UXOs often have to be “destroyed right there,” as noted by Anna Bouchier, Swiss director at Anti-personnel Mine Demining Product Development (APOPO). This poses particular challenges in agricultural regions, where land is desperately needed for cultivation. In many areas, fields are already being used for agriculture despite not being entirely mine-free.

These discussions helped identify four key areas where our solution could offer substantial benefits: time efficiency, cost reduction, environmental sustainability, and improved safety for human operators. Stakeholders also helped define the contexts in which our approach would be most impactful. For instance, NGOs operating in humid environments frequently encounter landmines with cracked or degraded casings, making them especially suitable for bacterial infiltration and degradation.

Beyond traditional landmines, our approach could also be extended to target other UXOs such as Improvised Explosive Devices (IEDs) and mortar rounds. These often involve lightweight plastic casings or partial failures on impact, leaving them cracked and leaking TNT. As Mark Wilkinson, humanitarian response team leader at DanChurchAid (DCA), explained: “A lot of them are cracked and broken with TNT inside.”

Because a 100% clearance rate is essential in demining operations, any novel approach must complement and integrate with established methods. Stakeholders emphasized that our solution’s strengths lie in its deployability, speed, and relative affordability. As Bill Morse (CSHD) put it: “Anything that’s going to reduce the number of landmines is going to be… a wonderful addition to what we have.”

Biosafety emerged as a major consideration due to the environmental release of genetically modified organisms (GMOs). Stakeholders appreciated the inclusion of kill-switch mechanisms but highlighted the importance of contingency measures in case of failure— for example, mutations that disable key regulatory elements or promoters. Concerns were also raised about the potential for horizontal gene transfer (HGT) between engineered bacteria and native soil microbes, which could facilitate the spread of antibiotic resistance genes (ARGs).

To mitigate these risks, we propose additional safety layers such as amino acid auxotrophy to restrict bacterial growth outside targeted environments—a strategy supported by prior studies [14]. Furthermore, we advocate for the use of the *Uplasmid* system developed by the 2023 iGEM EPFL team [33], which is specifically designed to minimize the risk of HGT.

These stakeholder insights were instrumental in refining both the technical aspects and the implementation strategy of our project, ensuring that our approach aligns with real-world needs and constraints in the field of humanitarian demining.

## Discussion

Our results demonstrate the feasibility and potential of using engineered bacteria and drone-based delivery systems for safe, targeted, and environmentally sustainable landmine and UXO remediation. Our colorimetric assay yielded particularly promising results. In sample 1 (Fig. 6b), the violet dye was visible, indicating the presence of TNT. In contrast, sample 2 (Fig. 6b) showed no violet dye, confirming TNT degradation. The assay reliably demonstrated that a decrease in the dye’s intensity correlates with reduced TNT concentration, highlighting its sensitivity and ability to distinguish between different TNT levels (Fig. 6a). Similarly, native *E. coli* exposed to 30 ng/µL of TNT showed no purple dye (Fig. 6c), consistent with the natural expression of *nfsA, nfsB*, and *nemA* in *E. coli* BL21 [4].

Additionally, upon exposure to TNT, our engineered bacteria formed a dark orange product, likely a Meisenheimer complex. This complex, associated with reduction of TNT’s aromatic ring and nitrate release, further suggests effective TNT degradation (Fig. 7).

Furthermore, our LC-MS analysis confirmed that TNT concentration in control samples (without bacteria) remained stable, indicating no spontaneous degradation or evaporation during the experiment (Fig. 8). Interestingly, both engineered and wild-type strains exhibited similar TNT degradation capabilities. When exposed to an initial concentration of 70 µg/mL of TNT, we observed an immediate decrease in concentration, followed by a transient increase and then another decline. This pattern suggests that bacteria initially sequester TNT before initiating its degradation, which may account for discrepancies between the initial input and early measured concentrations.

To improve degradation efficiency, we replaced the TNT-inducible *yqjF3rd* promoter with the constitutive *J23100* promoter. Constitutive expression of *nfsA* significantly enhanced TNT breakdown during the early phase (0–2 hours) compared to both the original engineered system and native *E. coli* (Fig. 8). These findings indicate that constitutive expression of *nfsA* offers a more rapid and efficient degradation strategy.

Our TNT fluorometric assay revealed a delayed onset of fluorescence, suggesting that the *yqjF3rd* promoter is not activated by TNT itself, but by one of its degradation intermediates. Based on prior work by the 2020 iGEM NEFU team [2], we hypothesize that the *yqjF3rd* promoter, optimized for DNT detection (a TNT degradation product), may have reduced sensitivity to TNT. This could explain both the observed fluorescence delay and the comparable degradation rates between engineered and unmodified strains.

Future work will focus on optimizing bacterial degradation of TNT, particularly using compacted TNT to better mimic conditions inside a landmine. Further efforts should also target additional explosives such as RDX and PETN. Although we have designed constructs for RDX degradation, they have yet to be tested experimentally.

Ensuring biosafety remains a critical concern, especially as we move toward field applications. Key safety features, such as the genetic AND gate and kill switch, have not yet been experimentally validated. It will also be important to address degradation of potentially toxic byproducts like formaldehyde to minimize environmental impact.

## Conclusion

Here, we evaluate the feasibility of an integrated synthetic biology and robotics approach for the clearance of landmines and UXOs. Our findings highlight three main areas of promise. First, our modelling work showed that TNT/DNT-induced luminescent bacteria could detect UXOs within a 5 cm radius under field conditions. Second, we developed a low-cost, functional drone prototype capable of detecting UXOs and delivering engineered bacteria or enzymes directly onto target sites. This prototype represents an accessible, low-cost alternative to high-end commercial drones. Third, we demonstrated effective TNT degradation using bacteria that constitutively express the *nfsA* enzyme, achieving the breakdown of 140 µg of TNT (at 70 µg/mL) in approximately 700 minutes.

Extensive interviews with demining stakeholders helped assess the real-world relevance of our approach. The feedback we received was largely positive, with many emphasizing the potential of our system to complement existing demining methods.

Overall, our work lays the foundation for integrating synthetic biology into practical, scalable mine-clearance strategies. Together, these advances highlight the promise of synthetic biology as a transformative tool for humanitarian demining, paving the way for safer, faster, and more accessible clearance of explosive remnants of war.

## Acknowledgment

We would like to express our gratitude to Philippe Abdel Sayed, our laboratory safety o”cer, for providing access to laboratory facilities. Our appreciation also goes to the Central Environmental Laboratory (CEL) at EPFL for conducting our LC-MS measurements, with particular thanks to Dominique Grandjean and Florian Breider. We are grateful to Armasuisse for supplying us with TNT for our research. Finally, we would like to thank the iGEM community for their inspiration, feedback, and support throughout this project.

## Ethics Statement

This study was conducted in accordance with the ethical guidelines from EPFL and the iGEM community for synthetic biology research.

## Data Availability Statement

All data and genetic constructs developed in this study are available through the iGEM Registry of Standard Biological Parts and the Synplode team iGEM wiki (Synplode).

## Supplementary Material

### Stakeholder engagement

During our study, we interviewed the following stakeholders:

#### NGOs

- Irene Rohner, Managing Co-Director of Welt Ohne Minen (World without mines), a Swiss foundation dedicated to supporting the eradication of landmines globally through fundraising initiatives.
- Representative of The Geneva International Centre for Humanitarian Demining (GICHD), a non profit organisation based in Geneva, Switzerland, dedicated to reducing the impact of landmines and explosive remnants of war (ERW).
- Bill Morse, chairman of Cambodian Self Help Demining (CSHD), a non-profit organisation focused on clearing landmines and UXOs in rural, low-priority Cambodian villages.
- Mark Wilkinson, Chief Technical Advisor of DanChurchAid (DCA), a Danish humanitarian organisation focused on fighting poverty and providing emergency relief in conflict- and disaster-affected regions.
- Dr Ahmed Al Zubaidi, co-founder of The Health and Social Care Organization in Iraq (IHSCO), a nonprofit humanitarian organisation focused on improving the health, social welfare, and quality of life for vulnerable populations in Iraq.
- Anna Bouchier, Swiss director of Anti-personnel Mine Demining Product Development (APOPO), a Belgian NGO that trains animals, particularly rats, to detect landmines and tuberculosis. sustainable development, humanitarian action, and peace.

#### Defence specialists

- Gäumann Jonathan, Senior Innovator and Security & Defence Specialist of RUAG, a Swiss technology company with a deep partnership with the Swiss Army, specialising in the development and provision of advanced defence technology solutions.
- Colonel Alexandro Spora, Chief of Deminers from Armasuisse.

#### Public organisations

- Dr. Anne-Laure Gasser, an employee in the science and technology division of Armasuisse, the procurement and technology centre for the Swiss Federal Department of Defence, Civil Protection, and Sport (DDPS).
- Tjark Thiele, employee of the United Nations Mine Action Service (UNMAS), a branch of the UN responsible for coordinating and implementing mine action initiatives in areas affected by landmines, UXOs, and improvised explosive devices (IEDs).

#### Academic researchers

- Dr. Jed. O. Eberly, PhD, associate professor at Montana State University responsible for developing a microbial sensor for RDX by engineering novel translational riboswitches (Eberly et al., 2019).
- Dr. Mariazel Maqueda-López, Head of the PeaceTech Division at the EssentialTech Centre of EPFL, dedicated to harnessing science and technology to support sustainable development, humanitarian action, and peace.

## Footnotes

1 SYNPLODE was part of the 2024 International Genetically Engineered Machine (iGEM) competition.

2 All sequences were codon-optimised for *E. coli* using GenSmart (GenScript) Optimization tool, and synthe-sised by GenScript, IDT or Twist Biosciences.

## References

[1] GICHD. A Study of Manual Mine Clearance - History, Summary and Conclusions of a Study of Manual Mine Clearance - World | ReliefWeb. Aug. 1, 2005. URL:https://reliefweb.int/report/world/study-manual-mine-clearance-history-summary-and-conclusions-study-manual-mine-clearance (visited on 11/18/2024).

[2] Team:NEFU China/Parts - 2020.igem.org. 2020. URL: https://2020.igem.org/Team:NEFU_China/Parts (visited on 11/10/2024).

[3] Benjamin Shemer et al. “Detection of buried explosives with immobilized bacterial bioreporters”. In: Microbial Biotechnology 14.1 (2021). Number: 1 _eprint: https://onlinelibrary.wiley.com/doi/pdf/10.1111/1751-7915.13683, pp. 251–261. ISSN: 1751-7915. DOI: 10.1111/1751-7915.13683. URL: https://onlinelibrary.wiley.com/doi/abs/10.1111/1751-7915.13683 (visited on 11/10/2024).

[4] M. Mar González-Pérez et al. “Escherichia coli has multiple enzymes that attack TNT and release nitrogen for growth”. In: Environmental Microbiology 9.6 (June 2007). Number: 6, pp. 1535–1540. ISSN: 1462-2912. DOI: 10.1111/j.1462-2920.2007.01272.x.

[5] Helena M. B. Seth-Smith et al. “Cloning, sequencing, and characterization of the hexahydro-1,3,5-Trinitro-1,3,5-triazine degradation gene cluster from Rhodococcus rhodochrous”. In: Applied and Environmental Microbiology 68.10 (Oct. 2002). Number: 10, pp. 4764–4771. ISSN: 0099-2240. DOI: 10.1128/AEM.68.10.4764-4771.2002.

[6] MM Ederer, TA Lewis, and RL Crawford. “2,4,6-Trinitrotoluene (TNT) transformation by clostridia isolated from a munition-fed bioreactor: comparison with non-adapted bacteria”. In: Journal of Industrial Microbiology and Biotechnology 18.2 (Feb. 1, 1997). Number: 2-3, pp. 82–88. ISSN: 1367-5435. DOI: 10.1038/sj.jim.2900257. URL: https://doi.org/10.1038/sj.jim.2900257 (visited on 11/10/2024).

[7] NG McCormick, FE Feeherry, and HS Levinson. “Microbial transformation of 2,4,6-trinitrotoluene and other nitroaromatic compounds”. In: Applied and Environmental Microbiology 31.6 (June 1976). Number: 6 Publisher: American Society for Microbiology, pp. 949–958. DOI:10.1128/aem.31.6.949-958.1976. URL: https://journals.asm.org/doi/abs/10.1128/aem.31.6.949-958.1976 (visited on 11/10/2024).

[8] M. E. Fuller and J. F. Manning. “Aerobic gram-positive and gram-negative bacteria exhibit di!erential sensitivity to and transformation of 2,4,6-trinitrotoluene (TNT)”. In: Current Microbiology 35.2 (Aug. 1997). Number: 2, pp. 77–83. ISSN: 0343-8651. DOI: 10.1007/s002849900216.

[9] Muratnal. “Effective Degradation of 2,4,6-Trinitrotoluene (TNT) with a Bacterial Consortium Developed from High TNT-degrading Bacteria Isolated from TNT-contaminated Soil”. In: Hacettepe Journal of Biology and Chemistry 3.46 (Oct. 18, 2018). Number: 46, pp. 445–455. ISSN: 1303-5002. DOI: 10.15671/HJBC.2018.252.URL:chem-volume-46-issue-3-pg-445-455.pdf (visited on 11/10/2024).

[10] Richard Williams et al. “Biotransformation of Explosives by the Old Yellow Enzyme Family of Flavoproteins”. In: Applied and environmental microbiology 70 (June 7, 2004), pp. 3566–74. DOI: 10.1128/AEM.70.6.3566-3574.2004.

[11] Diane Fournier et al. “Determination of key metabolites during biodegradation of hexahydro-1,3,5trinitro-1,3,5-triazine with Rhodococcus sp. strain DN22”. In: Applied and Environmental Microbiology 68.1 (Jan. 2002), pp. 166–172. ISSN: 0099-2240. DOI: 10.1128/AEM.68.1.166-172.2002.

[12] Tae Seok Moon et al. “Genetic programs constructed from layered logic gates in single cells”. In: Nature 491.7423 (Nov. 2012). Number: 7423 Publisher: Nature Publishing Group, pp. 249–253. ISSN: 1476-4687. DOI: 10.1038/nature11516. URL: https://www.nature.com/articles/nature11516 (visited on 11/10/2024).

[13] P. Bernard and M. Couturier. “Cell killing by the F plasmid CcdB protein involves poisoning of DNA-topoisomerase II complexes”. In: Journal of Molecular Biology 226.3 (Aug. 5, 1992). Number: 3, pp. 735–745. ISSN: 0022-2836. DOI: 10.1016/0022-2836(92)90629-x.

[14] Christopher M. Whitford et al. “Auxotrophy to Xeno-DNA: an exploration of combinatorial mechanisms for a high-fidelity biosafety system for synthetic biology applications”. In: Journal of Biological Engineering 12 (Aug. 14, 2018), p. 13. DOI: 10.1186/s13036-018-0105-8. URL: https://pmc.ncbi.nlm.nih.gov/articles/PMC6090650/ (visited on 11/10/2024).

[15] Center for International Stabilization {and} Recovery (CISR). Study of the E!ects of Aging on Landmines. James Madison University, 2010. URL: https://www.nature.com/articles/nature11516

[16] R. Bello. “Literature Review on Landmines and Detection Methods”. In: 2013. URL:semanticscholar.org/paper/Literature-Review-on-Landmines-and-Detection-Bello/af41c436254102691ced90701e22 (visited on 11/13/2024).

[17] Nguyen Trung Thành, Dinh Nho Hào, and Hichem Sahli. “Infrared Thermography for Land Mine Detection”. In: Augmented Vision Perception in Infrared: Algorithms and Applied Systems. Ed. by Riad I. Hammoud. London: Springer London, 2009, pp. 3–36. ş%)&: 978-1-84800-277-7. DOI: 10.1007/978-1-84800-277-7_1. URL: https://doi.org/10.1007/978-1-84800-277-7_1.

[18] Theo John Marshall, John Winspere Holmes, and Calvin Rose. Soil Physics. Cambridge University Press, 1996.

[19] Etai Shpigel et al. “Introduction of quorum sensing elements into bacterial bioreporter circuits enhances explosives’ detection capabilities”. In: Engineering in Life Sciences 22.3 (2022). Number: 3-4 _eprint: https://onlinelibrary.wiley.com/doi/pdf/10.1002/elsc.202100134, pp. 308–318. ISSN: 1618-2863. DOI: 10.1002/elsc.202100134. URL::https://onlinelibrary.wiley.com/doi/abs/10.1002/elsc.202100134 (visited on 11/10/2024).

[20] Sharon Yagur-Kroll et al. “Escherichia coli bioreporters for the detection of 2,4-dinitrotoluene and 2,4,6-trinitrotoluene”. In: Applied Microbiology and Biotechnology 98.2 (Jan. 1, 2014). Number: 2, pp. 885–895.ISSN: 1432-0614. DOI: 10.1007/s00253-013-4888-8. URL:https://doi.org/10.1007/s00253-013-4888-8 (visited on 11/10/2024).

[21] Benjamin Shemer et al. “Aerobic Transformation of 2,4-Dinitrotoluene by Escherichia coli and Its Implications for the Detection of Trace Explosives”. In: Applied and Environmental Microbiology 84.4 (Feb. 15, 2018). Number: 4, e01729–17. ISSN: 1098-5336. DOI: 10.1128/AEM.01729-17.

[22] Stavrini Papadaki et al. “Dual-expression system for blue fluorescent protein optimization”. In: Scientific Reports 12.1 (June 17, 2022). Publisher: Nature Publishing Group, p. 10190. ISSN: 2045-2322. DOI: 10.1038/s41598-022-13214-0. URL: https://www.nature.com/articles/s41598-022-13214-0 (visited on 12/06/2024).

[23] Jed O. Eberly et al. “Detection of hexahydro-1,3-5-trinitro-1,3,5-triazine (RDX) with a microbial sensor”. In: The Journal of General and Applied Microbiology 65.3 (July 19, 2019). Number: 3, pp. 145–150. ISSN: 1349-8037. DOI: 10.2323/jgam.2018.08.001.

[24] Ali Seleit, Alexander Aulehla, and Alexandre Paix. “Endogenous protein tagging in medaka using a simplified CRISPR/Cas9 knock-in approach”. In: eLife 10 (Dec. 6, 2021), e75050. ISSN: 2050-084X. DOI: 10.7554/eLife.75050. URL:https://elifesciences.org/articles/75050 (visited on 11/14/2024).

[25] Katie J. Denby et al. “The mechanism of a formaldehyde-sensing transcriptional regulator”. In: Scientific Reports 6.1 (Dec. 9, 2016). Number: 1 Publisher: Nature Publishing Group, p. 38879. ISSN: 2045-2322. DOI: 10.1038/srep38879. URL: https://www.nature.com/articles/srep38879 (visited on 11/10/2024).

[26] Ankith Sharma, Rajdeep Chowdhury, and Siegfried M. Musser. “Oligomerization state of the functional bacterial twin-arginine translocation (Tat) receptor complex”. In: Communications Biology 5.1 (Sept. 19, 2022), p. 988. ISSN: 2399-3642. DOI: 10.1038/s42003-022-03952-2. URL:https://www.nature.com/articles/s42003-022-03952-2 (visited on 11/14/2024).

[27] Frank Schnütgen et al. “A directional strategy for monitoring Cre-mediated recombination at the cellular level in the mouse”. In: Nature Biotechnology 21.5 (May 2003). Number: 5 Publisher: Nature Publishing Group, pp. 562–565. ISSN: 1546-1696. DOI: 10.1038/nbt811. URL:https://www.nature.com/articles/nbt811 (visited on 11/10/2024).

[28] Shana Topp et al. “Synthetic riboswitches that induce gene expression in diverse bacterial species”. In: Applied and Environmental Microbiology 76.23 (Dec. 2010). Number: 23, pp. 7881–7884. ISSN: 1098-5336. DOI: 10.1128/AEM.01537-10.

[29] Ayşem Üzer, Erol Erça%, and Reşat Apak. “Selective spectrophotometric determination of TNT in soil and water with dicyclohexylamine extraction”. In: Analytica Chimica Acta 534.2 (Apr. 8, 2005), pp. 307–317.URL: https://www.sciencedirect.com/science/article/pii/S0003267004015739 (visited on 11/18/2024).

[30] Bibiana Báez et al. “Detection of chemical signatures from TNT buried in sand at various ambient conditions: phase II”. In: Detection and Remediation Technologies for Mines and Minelike Targets XI. Detection and Remediation Technologies for Mines and Minelike Targets XI. Vol. 6217. SPIE, May 18, 2006, pp. 495–504. ISSN: 10.1117/12.666012. URL: https://www.spiedigitallibrary.org/conference-proceedingsof-spie/6217/62171M/Detection-of-chemical-signatures-from-TNT-buried-in-sand-at/10.1117/12. 666012.full (visited on 11/18/2024).

[31] Tal Elad et al. “Enhancing DNT Detection by a Bacterial Bioreporter: Directed Evolution of the Tran-scriptional Activator YhaJ”. In: Frontiers in Bioengineering and Biotechnology 10 (2022). ISSN: 2296-4185. DOI: 10.3389/fbioe.2022.821835.URL:https://www.frontiersin.org/journals/bioengineering-and-biotechnology/articles/10.3389/fbioe.2022.821835.

[32] Justin Merritt, Hidenobu Senpuku, and Jens Kreth. “Let there be bioluminescence: development of a biophotonic imaging platform for in situ analyses of oral biofilms in animal models”. In: Environmental Microbiology 18.1 (Jan. 2016). Number: 1, pp. 174–190. ISSN: 1462-2920. DOI: 10.1111/1462-2920.12953.

[33] EPFL iGEM 2023 - 48C. URL: https://2023.igem.wiki/epfl/safety#designs (visited on 12/06/2024).

